# From adverse to beneficial – contrasting dietary effects of freshwater mixotrophs on zooplankton

**DOI:** 10.1101/2021.01.08.425733

**Authors:** Csaba F. Vad, Claudia Schneider, Robert Fischer, Martin J. Kainz, Robert Ptacnik

## Abstract

1. The importance of mixotrophic algae as key bacterivores in microbial food webs is increasingly acknowledged, but their effects on the next trophic level remain poorly understood. Their high stoichiometric food quality is contrasted by anti-grazing strategies.
2. We tested the quality of freshwater mixotrophs as prey for zooplankton, using four non-colonial chrysophyte species and a cryptophyte as a high quality reference food. We (1) analyzed the stoichiometric and biochemical (fatty acid) composition of the mixotrophs, and (2) quantified their dietary effects on *Daphnia longispina* survival.
3. Survival of *D. longispina* significantly depended on the identity of species provided as food, ranging from higher to lower as compared to starvation. This was not reflected in differences in cellular stoichiometry or fatty acid profiles of the mixotrophs. We suggest that toxicity may be the driver for the observed differences.
4. Generalization of the dietary effects of mixotrophic chrysophytes does not appear straightforward. Besides fundamental species-specific differences, potential toxic effects may vary depending on environmental cues or physiological strategies. Notably in our study, *Ochromonas tuberculata*, a species previously reported to be deleterious turned out to be a beneficial food source, in terms of enabling high survival of *D. longispina*.
5. We challenge the generality of the assumption that chrysophytes are of low value as food for zooplankton. We recommend that future studies test how environmental conditions and physiological strategies shape the quality of mixotrophs as food for consumers at higher trophic levels, specifically focusing on effects of dietary toxicity.

## 1. Introduction

Mixotrophic protists are common in the plankton of freshwater and marine ecosystems, and can represent a dominant component of phytoplankton in the surface waters of oceans (Zubkov & Tarran, 2008; Hartmann *et al*., 2012; Leles *et al*., 2019) and lakes (Watson, McCauley & Downing, 1997; Pålsson & Granéli, 2004; Ptacnik *et al*., 2008). Their success appears to be driven by their ability to combine photosynthesis with the ingestion of small particles, e.g. bacteria-sized plankton, which can give them a competitive edge over specialized phototrophic and heterotrophic competitors, particularly under nutrient limitation (Katechakis & Stibor, 2006; Kamjunke, Henrichs & Gaedke, 2007; Fischer *et al*., 2017). Mixotrophs often account for the bulk of bacterivory in near-surface waters (Bird & Kalff, 1987; Isaksson *et al*., 1999; Domaizon, Viboud & Fontvieille, 2003; Unrein *et al*., 2014), and thereby may play a key role for carbon cycling at the base of the aquatic food web (Mitra *et al*., 2014). Despite the increasing interest in understanding their trophic function as consumers, i.e., quantifying their importance as bacterivores (Hartmann *et al*., 2012; Ptacnik *et al*., 2016; Anderson, Jürgens & Hansen, 2017; Beisner, Grossart & Gasol, 2019), little is known about their role as prey for zooplankton. As a consequence, their impact on consumers at higher trophic levels remains unclear.

Recent global models on the importance of mixotrophic plankton suggest that mixotrophy increases trophic transfer efficiency in the pelagic food web (Ward & Follows, 2016; Stoecker *et al*., 2017). This is in line with the observation that mixotrophs have lower and less variable cellular ratios of carbon to nutrients (i.e., C:P and C:N) than strict photoautotrophs (Katechakis *et al*., 2005; Moorthi *et al*., 2017), which may reduce the elemental mismatch of consumers’ nutrient demands and the nutrient balance of their algal prey. A stoichiometric mismatch results in nutrient limitation and can constrain consumer growth (Sterner & Elser, 2002), therefore, its reduction via an increased proportion of mixotrophs in phytoplankton may have a stabilizing effect on secondary production (Moorthi *et al*., 2017). At the same time, most harmful (eukaryotic) algal blooms, with detrimental effects on secondary production, are caused by mixotrophic species (Burkholder, Glibert & Skelton, 2008; Watson *et al*., 2015; Flynn *et al*., 2018), suggesting a link between mixotrophy and toxicity. Whether this contrasting evidence on the food quality of mixotrophs are specific to major taxonomic groups or are related to inter-specific differences, is an important question to be solved to better understand the functional roles of mixotrophs in both microbial and associated food webs.

In lake phytoplankton, there are two widespread groups of nanoflagellates (<20 μm) in which mixotrophy is prevalent: chrysophytes and cryptophytes (Sanders & Porter, 1988). Among these groups, cryptophytes are generally acknowledged as high-quality food for zooplankton based on their high cellular content of essential fatty acids (Ahlgren *et al*., 1990; Brett & Müller-Navarra, 1997; von Elert & Stampfl, 2000). Much less is known about the dietary effects of chrysophytes on higher trophic level, and current reports are rather inconsistent. For example, empirical observations suggest that mixotrophic chrysophytes may account for the bulk of zooplankton diet in some freshwater systems (Bertoni, Callieri & Corno, 2002). In contrast, colony-forming species, such as the widespread *Dinobryon*, are largely grazing resistant and of low diet quality due to poorly digestible cellulose structures (Vad *et al*., 2020). In addition, experimental studies on non-colony-forming chrysophytes suggest toxic effects on zooplankton growth, reproduction and survival (Table 1). These experiments are, however, limited to a rather narrow set of species (i.e., 6 taxa belonging to 3 genera) and, to the best of our knowledge, no study tested the effect of more than one chrysophyte species simultaneously, i.e., under the same conditions, in feeding experiments with zooplankton.

**Table 1.**
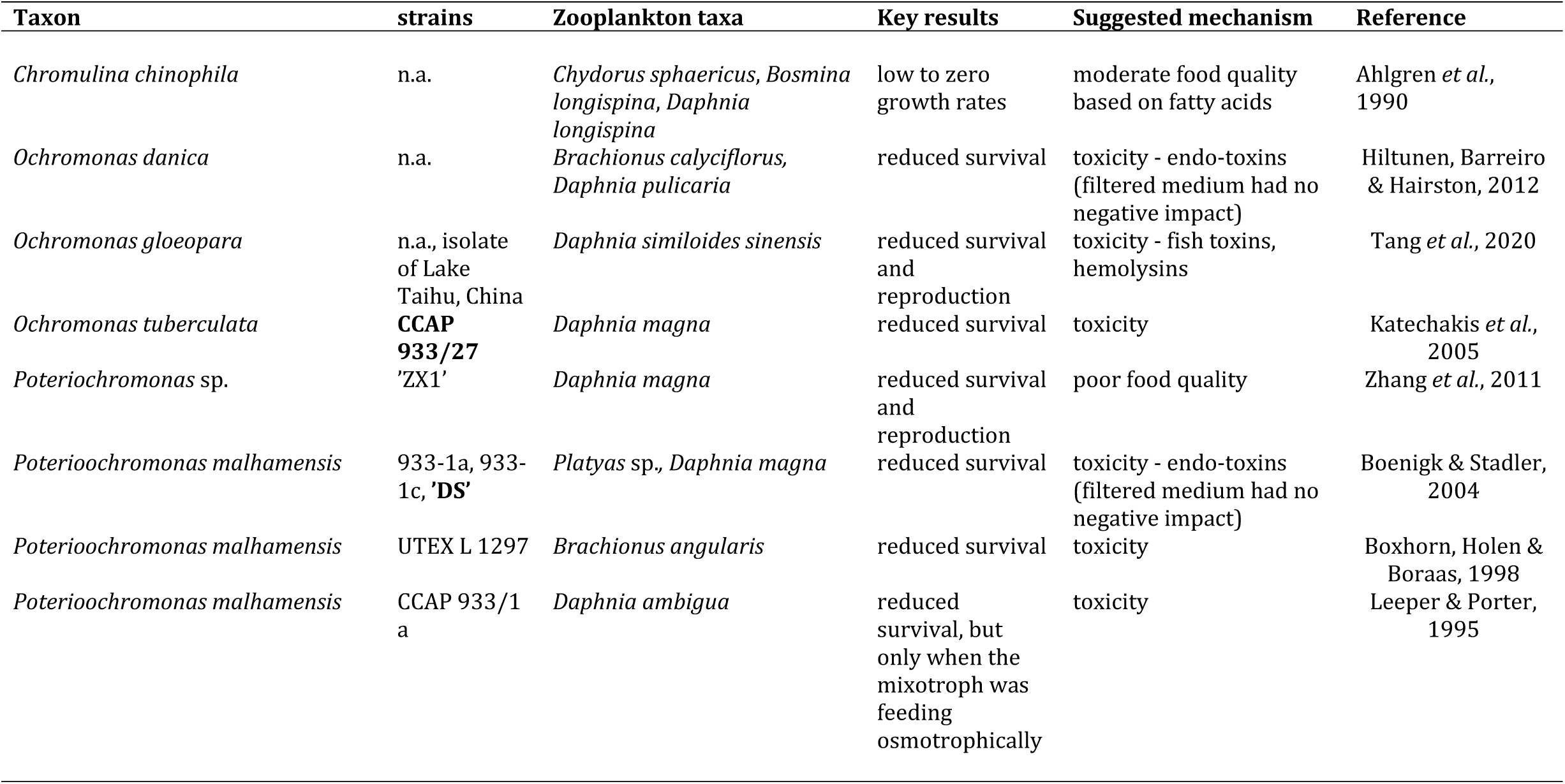
A review of feeding experiments performed with non-colony-forming mixotrophic chrysophytes and zooplankton. Strains used in the present study are highlighted by bold letters. Note that while toxic effects are regularly reported, identification of toxins was only performed by Tang *et al*. (2020).

The biochemical food quality of chrysophytes represents a further knowledge gap as comprehensive studies on their fatty acid profiles are largely lacking (but see Ahlgren *et al*., 1990; Boëchat *et al*., 2007). Overall, there appear to be fundamental differences in the dietary quality of freshwater mixotrophs. In contrast to cryptophytes, chrysophytes generally are assumed to represent low diet quality, and can be even toxic food for consumers (Table 1). This difference, apparently specific to major groups may provide an explanation for the contradiction in the profitability of mixotrophs as food sources. However, multi-species feeding experiments are still needed to be able to generalize this observation.

Here we test the effect of a diet consisting of non-colony forming chrysophytes species on the cladoceran zooplankter *Daphnia*. As a large filter-feeder crustacean, it represent a key element in lakes by contributing to the structuring of food webs by linking primary production with higher trophic levels (Carpenter & Kitchell, 1996; Sommer & Stibor, 2002). There is ample evidence that the efficiency of trophic transfer through zooplankton depends strongly on algal food quantity and food quality (Sterner & Hessen, 1994; Müller-Navarra & Lampert, 1996; Brett & Müller-Navarra, 1997; Sterner & Schulz, 1998). Assuming predominantly detrimental dietary effects of mixotrophic chrysophytes on consumers, as suggested by previous experimental studies (Table 1), a shift in the phytoplankton assemblage towards this taxonomic group may have profound effects on zooplankton secondary production and consequently higher trophic levels (e.g. fish).

The specific aims of this study are to, a) identify the stoichiometric (molar C:N:P ratios) and biochemical (fatty acids) compositions of multiple non-colony-forming chrysophyte species, and, b) perform a comparative test to quantify their nutritional adequacy and potential toxicity to *Daphnia*. Based on the available evidence, we hypothesize that chrysophytes will represent lower-quality food sources, being in general detrimental compared to *Cryptomonas*, used as a high-quality reference food.

## 2. Material and methods

### 2.1. Study species and cultivation conditions

In the study we used five algal species, including four chrysophytes species, *Ochromonas danica* (strain: SAG 933-7), *Ochromonas tuberculata* (strain: CCAP 933/27), *Ochromonas* sp. (own isolate from Lake Lunz; available through the AquaScale lab), *Poterioochromonas malhamensis* (strain DS, formerly used under the names ‘*Ochromonas* sp.’, ‘*Ochromonas* sp. strain DS’ and ‘*Ochromonas* DS’, e.g. Hahn & Höfle, 1998; Boenigk & Stadler, 2004; Boenigk *et al*., 2004), and one cryptophyte species, *Cryptomonas* sp. (strain: SAG 26.80). Note that *Ochromonas* sp. was not used in experiment 1 (‘stoichiometric and biochemical composition’), because it was only isolated from Lake Lunz (situated in the Eastern Alps in Austria, 608 m a.s.l, 47°51.2′N 15°3.1′E) after this first experiment was conducted. All algae cultures were non-axenic, i.e., cultures contained not identified heterotrophic bacteria. Cultures were cultivated in a walk-in environmental chamber at a constant temperature of 18°C, a 16:8 hours light:dark photoperiod, and saturating irradiance of approximately 170 μmol photon m^− 2^ s^− 1^. Prior to the experiments, algae cultures were pre-cultivated for several weeks using a medium consisting of 90% sterile-filtered lake water (0.2-μm pore size) from the oligotrophic Lake Lunz (5–8 μg total phosphorus L^-1^) enriched with WEES medium (10% of total volume) modified by leaving out soil extract (WEES Medium Recipe vers. 03.2007, Göttingen University, Kies, 1967). Regular dilution of the cultures ensured to keep cultures in exponential growth prior to the experiment (cell densities were monitored by flow cytometry).

For experiment 2 (‘feeding experiment’), algal biomass was based on cell concentrations and species-specific cell biovolumes. Algal biovolumes (Table 1) were calculated using conversion factors for corresponding geometrical shapes of each respective species (Hillebrand *et al*., 1999). To this end, axial dimensions of at least 30 cells per species were microscopically measured (AxioVision Software, Carl Zeiss AG, Jena, Germany). Carbon biomass was estimated from biovolume applying a conversion factor of 14% of algal wet weight (Vadstein *et al*., 1988).

The zooplankton species used in this study was the cladoceran *Daphnia cf. longispina* (belonging to the *D. longispina* complex, hereinafter referred to as *D. longispina*). Prior to the experiment, a *D. longispina* clone was isolated from Lake Lunz and cultures were kept under the same temperature and light conditions as described for the algae, on sterile-filtered lake water. They were fed with *Cryptomonas* sp., which is known to be of high nutritional value for zooplankton (Martin-Creuzburg, Elert & Hoffmann, 2008). Though we did not test in the specific strain we used, it is a representative of a genus where mixotrophy is widespread (Tranvik, Porter & Sieburth, 1989; Domaizon *et al*., 2003). For the ‘feeding experiment’, third-brood neonates (<24 hours old) from a previously established F2 generation were used. This procedure ensured a standardised response of juveniles during the experiment and a reduction of maternal effects.

### 2.2. Experiment 1: Stoichiometric and biochemical composition

In this first experiment we assessed and compared the cellular stoichiometry and fatty acid composition of *O. danica, O. tuberculata*., *P*.*malhamensis* and *Cryptomonas* sp., as indicators of their food quality. Algae were either grown predominately photoautotrophically or mixotrophically, respectively. To this end, species were grown individually in Erlenmeyer flasks (200 mL culture volume; three replicates per species) under similar conditions as described for the pre-cultivation, however, we further modified the growth media by only using 10% of the phosphorus concentration relative to the original recipe (final P concentration in media: approx. 50 μg L^-1^). This ensured to expose algae to conditions more similar to their natural environment where chrysophytes are often dominant under P-limited conditions (Ptacnik *et al*., 2008; van Donk *et al*., 2009). We assumed that bacteria present in the culture were energy-limited, i.e., besides low levels of background dissolved organic carbon (DOC) from the lake water, no C-sources were present in the media, and hence bacteria abundances would not be sufficient to support mixotrophic growth. Under these conditions we assumed predominately photoautotrophic nutrition by the tested species.

To induce predominantly mixotrophic nutrition, i.e., elevated rates of bacterivory, bacterial growth was stimulated in another batch of Erlenmeyer flasks (three replicates per species) by daily addition of an allochthonous carbon source (i.e., α-D-glucose monohydrate, Merck KGaA, Darmstadt, Germany, supplied at 1 mg L^-1^). We assumed that bacteria would be better competitors for the added glucose at these concentrations (Kamjunke *et al*., 2008), and hence exclude the possibility that glucose was directly used by the algae for growth. Indeed, osmotrophic uptake of dissolved organic substrates in phytoplankton is common, however, bacteria should generally have higher affinities for such substrates. Moreover, the underlying trade-offs of the mixotrophic nutrition suggest that mixotrophs may show only low uptake of dissolved substrates (Flynn & Mitra, 2009). Besides providing more natural conditions, we assumed that using a medium with low nutrient concentrations would also favour mixotrophic growth of the algae in the treatments with glucose addition. Many mixotrophic species only engage in, or have increased rates of bacterivory, when nutrients limit their growth (Nygaard & Tobiesen, 1993; Keller *et al*., 1994). For example, the mixotrophic chrysophyte *Uroglenopsis americana* expressed significantly lower rates of bacterivory when exposed to phosphorus levels >12.4 μg L^-1^ (Urabe, Gurung & Yoshida, 1999).

Cell densities were measured every three days using a flow cytometer (CytoFLEX, Beckman Coulter Life Sciences, Indianapolis, IN, USA). Manual gating was used to detect populations of *O. danica, O. tuberculata*., *P. malhamensis* and *Cryptomonas* sp., respectively (detection: 690 ± 50 nm versus 660 ± 20 nm). The experiment was terminated and samples for analyses were taken from cultures when they reached stationary phase, i.e., abundances stabilized over a few days (data not shown). In the treatments with predominately photoautotrophic growth, the tested chrysophytes did not show considerable growth resulting in insufficient biomass for further analyses (this was not the case with *Cryptomonas* sp.). We therefore only report stoichiometric and biochemical profiles of the tested species from the treatments with predominately mixotrophic growth.

Upon terminating the experiment, samples for particulate fractions (elemental stoichiometry and lipids and their fatty acids) were filtered on pre-combusted and acid washed glass microfiber filters (25 mm diameter, Whatman GF/F). Three separate filters were prepared per each experimental replicate (for C and N, for P, and for fatty acid analysis). Prior to measurements, C and N, and P filters were dried for at least 48 h in a drying chamber at 60°C and stored until analysis. Filters for fatty acid analysis were immediately frozen at -80°C.

For obtaining data on cellular stoichiometry, the amount of particulate organic carbon and nitrogen was determined by an elemental analyser (vario MICRO cube™, Elementar Analysensysteme GmbH, Germany). Particulate P was measured as orthophosphate by a molybdate reaction after sulfuric acid digestion (Solórzano & Sharp, 1980; Hansen & Koroleff, 1999).

The analysis of lipids and their fatty acids followed the protocol described by Heissenberger, Watzke & Kainz (2010). In brief, samples were freeze-dried and homogenized before analyses. Fatty acid methyl esters (FAME) were analysed by a gas chromatograph (TRACE GC Ultra™, Thermo Fisher Scientific, Waltham, MA, USA; equipped with a flame ionization detector) and separated with a Supelco™ SP-2560 column (Bellefonte, PA, USA). FAME were identified using known standards (37-component FAME Mix, Supelco 47885-U).

### 2.3. Experiment 2: Feeding experiment

We tested the effect of five different algal species, individually provided as food source, on survival and growth of *D. longispina*. This experiment consisted of six treatments, each representing a different algal food source: (1) *O. danica*, (2) *Ochromonas* sp., (3) *O. tuberculata*, (4) *P. malhamensis*, (5) *Cryptomonas* sp., used as a high-quality reference food and (6) a starvation treatment as control (Lampert, 1981). Each treatment was replicated five times. All tested algal species fall well in the size range (nano-flagellates) that can be exploited by *Daphnia*, therefore we neglected size-dependent differences in ingestion as driver of potentially observed dietary effects (Table 2).

**Table 2.** Mean (± SD) cell dimensions and biovolumes of all the algae used in the study. For species resembling a ‘prolate sphaeroid’ shape, biovolume calculations were based on length and width, while for spherical species it was approximated from cell diameter.

| Species | Length ($\mu\text{m}$) | Width ($\mu\text{m}$) | Diameter ($\mu\text{m}$) | Biovolume ($\mu\text{m}^3$) |
| --- | --- | --- | --- | --- |
| *Cryptomonas* sp. | $14.6 \pm 1.7$ | $8.3 \pm 0.9$ | - | $537.1 \pm 142.4$ |
| *Ochromonas danica* | $10.2 \pm 1.7$ | $7.8 \pm 1.4$ | - | $345.7 \pm 175.2$ |
| *Ochromonas tuberculata* | - | - | $11.1 \pm 2.1$ | $797.7 \pm 519.7$ |
| *Ochromonas* sp. | - | - | $6.9 \pm 0.6$ | $177.4 \pm 49.5$ |
| *Poterioochromonas malhamensis* | - | - | $7.2 \pm 1.1$ | $207.9 \pm 107.8$ |

At the start of the experiment, three *D. longispina* neonates per replicate were introduced to glass vials containing 30 mL of the experimental food suspension of the corresponding treatments. Each experimental food suspension consisted of sterile-filtered lake water, enriched with the respective algal species to attain food-saturated conditions of approximately 1 mg C L^-1^. The starvation treatment consisted of sterile-filtered lake water, without any food supply. Animals were transferred to fresh experimental food suspension every other day to maintain relatively stable food biomass (this was also done for the starvation treatment, to standardise handling). The experiment was run for 11 days (i.e., the day after *D. longispina* started releasing juveniles). The experimental temperature and light conditions were the same as described for the pre-cultivation. During the experiment, vials were gently mixed twice a day to reduce sedimentation and surface accumulation of the algae. Survival and the number of moults (i.e., the number of shed carapaces), as a proxy for growth (Hessen & Rukke, 2000), were recorded daily. Dead individuals were removed (animals were considered dead when lying on the bottom of the vial without any movement).

The four chrysophyte species were cultivated during the experiment as described for the pre-cultivation, however, the medium with reduced P concentration used in the first experiment was not sufficient to yield algal biomass required for the second experiment. For unknown reasons *Cryptomonas* sp. was insufficiently growing on the 10% WEES : 90% sterile-filtered lake water medium at the time of the second experiment, and we adjusted the proportion of WEES medium in experimental medium to 20%. We also stimulated bacterial growth (glucose addition; 1 mg L^-1^) to enhance mixotrophic growth, but only during pre-cultivation. This ensured that bacteria would not represent sufficient prey biomass to support survival and growth of *D. longispina*.

### 2.4. Data analysis

Temporal (daily) patterns of survival (i.e., percentage of living individuals) of *D. longispina* were analysed by testing for pairwise differences between each treatment and the starvation treatment via Wilcoxon rank-sum tests. Separate tests were run for each day and treatment. The use of a non-parametric test was needed as data were non-normally distributed in most cases.

As further indicators of the condition of animals, growth (cumulative number of moults) and the number of mature females of *D. longispina* at the termination of the experiment were also recorded. To test for significant differences among treatments, non-parametric Kruskal-Wallis tests were performed (because of non-normally distributed residuals of a previous one-way ANOVA, tested by Shapiro Wilk test). This was followed by Dunn’s *post hoc* test (p values were adjusted to account for multiple comparisons with the Holm method) to reveal pairwise differences. All statistical analyses were done with the R software (R Core Team, 2020).

## 3. Results

### 3.1. Stoichiometric and biochemical composition

The molar seston C:P ratios of all chrysophyte cultures ranged between 100 and 200, while N:P ranged between 20 and 30 (Table 3). In the *Cryptomonas* cultures, both ratios were dependent on whether it was grown with an external carbon source, with higher values when supplied with glucose.

**Table 3.** C:P and N:P elemental ratios of algae along with mass fractions (μg mg dry weight^-1^) of fatty acid groups, palmitoleic acid (16:1n-7), and the n-3/n-6 PUFA and algal/bacterial fatty acid ratios (data represent mean ± SD of three replicates). For chrysophytes, we present data resulting from the experiment with input of dissolved organic carbon (DOC), i.e., glucose, where they reached biomasses sufficient for the analysis. Fatty acid data on *Cryptomonas* with DOC-input are missing due to a technical error. (SAFA – saturated fatty acids; MUFA – monounsaturated fatty acids; PUFA – polyunsaturated fatty acids, used as indicators of mostly algae-produced fatty acids; BFA – bacterial fatty acids).

|  | *Cryptomonas*<br>sp. -DOC | *Cryptomonas*<br>sp. +DOC | *Ochromonas*<br>*danica* | *Ochromonas*<br>*tuberculata* | *Poterioochromonas*<br>*malhamensis* |
| --- | --- | --- | --- | --- | --- |
| C:P | 100.9 ± 12.7 | 244.1 ± 7.7 | 155.0 ± 27.1 | 156.4 ± 10.9 | 155.8 ± 21.0 |
| N:P | 16.7 ± 1.6 | 29.1 ± 1.2 | 22.5 ± 3.9 | 21.9 ± 0.7 | 21.3 ± 3.8 |
| SAFA | 27.3 ± 6.9 | - | 53.6 ± 9.2 | 39.8 ± 15.6 | 43.5 ± 6.2 |
| MUFA | 14.3 ± 3.4 | - | 16.3 ± 3.7 | 10.9 ± 4.2 | 14.7 ± 1.5 |
| PUFA | 31.5 ± 2.2 | - | 29.8 ± 7.7 | 6.9 ± 2.6 | 5.2 ± 0.6 |
| BFA | 11.0 ± 1.5 | - | 12.3 ± 2.8 | 7.9 ± 3.1 | 10.6 ± 1.9 |
| ω-3 PUFA | 23.3 ± 1.4 | - | 15.5 ± 4.3 | 3.9 ± 1.5 | 3.4 ± 0.2 |
| ω-6 PUFA | 8.1 ± 1.0 | - | 14.3 ± 3.4 | 2.9 ± 1.1 | 1.8 ± 0.5 |
| 16:1n-7 | 4.6 ± 2.1 | - | 1.3 ± 0.3 | 3.5 ± 1.4 | 0.6 ± 0.1 |
| n-3/n-6 | 2.9 ± 0.2 | - | 1.1 ± 0.1 | 1.3 ± 0.1 | 2.0 ± 0.5 |
| PUFA/BF |  |  |  |  |  |
| A | 2.9 ± 0.3 | - | 2.4 ± 0.1 | 0.9 ± 0.0 | 0.5 ± 0.0 |

Considering fatty acid profiles, *O. danica* was outstanding with a higher content of polyunsaturated fatty acids (PUFA) than *O. tuberculata* and *P. malhamensis*, at the same time comparable to that of *Cryptomonas* sp. (Table 3). Furthermore, all individual PUFA occurred in higher amounts in *O. danica* than in the other two chrysophytes (see Table S1 in Supporting Information).

### 3.2. Feeding experiment

Survival of *D. longispina* was significantly influenced by the food source provided. On *O. danica*, a severe drop could be observed already after one day, resulting in significantly lower survival than in the starvation treatment (Fig. 1). On *P. malhamensis*, high mortalities occurred later in the experiment leading to the death of all neonates by day 10 (significantly lower survival compared to the starvation treatment on days 8–10; Fig. 1). Survival on *Cryptomonas* sp., *Ochromonas* sp. and *O. tuberculata* stayed high throughout the experiment, and it was significantly higher compared to the starvation treatment towards the end of the experiment.

**Fig. 1.**
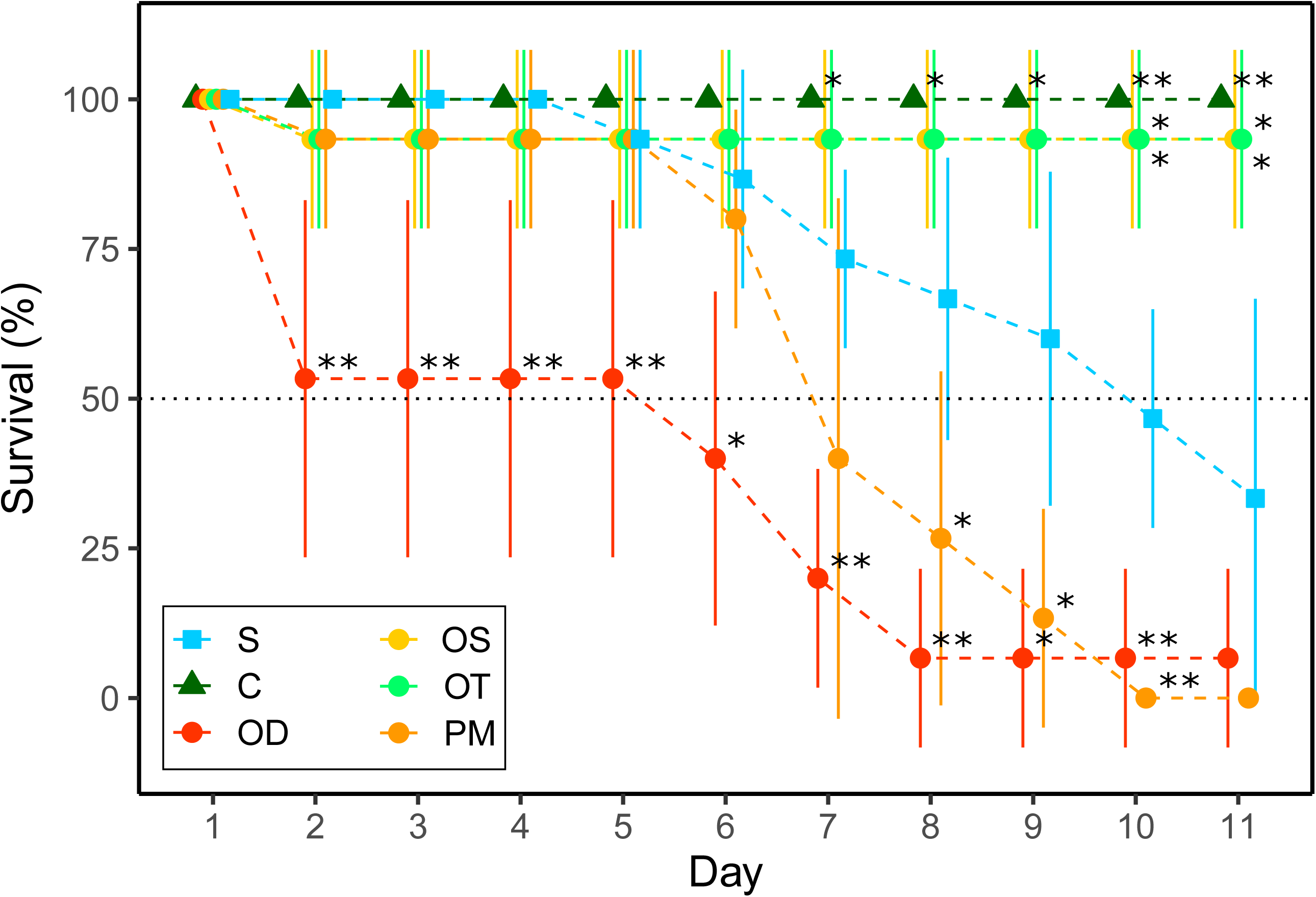
Temporal pattern in the percentage of surviving *D. longispina* (mean ± SD) on the different algae and the starvation treatment. Asterisks indicate significant differences in comparison to the starvation treatment based on pairwise Wilcoxon rank-sum tests (significance levels: p < 0.05 ‘*’, p < 0.01 ‘**’). Abbreviations: Starvation treatment (S), the reference food *Cryptomonas* sp. (C), *Ochromonas danica* (OD), *Ochromonas* sp.(OS), *Ochromonas tuberculata* (OT), *Poterioochromonas malhamensis* (PM).

The number of moults was also significantly influenced by the food source provided (Fig. 2, Kruskal-Wallis test: χ2 = 24.2, df = 5, p = < 0.001). Highest numbers of moults were observed on *Cryptomonas* sp., *Ochromonas* sp. and O. *tuberculata*, showing no statistical difference among these treatments. Moulting was suppressed on *O. danica* and *P. malhamensis*, and was significantly lower in these treatments compared to *Cryptomonas* sp., and was furthermore significantly reduced on *O. danica* compared to the other two *Ochromonas* species. Moulting rates in the starvation treatment were intermediate.

**Fig. 2.**
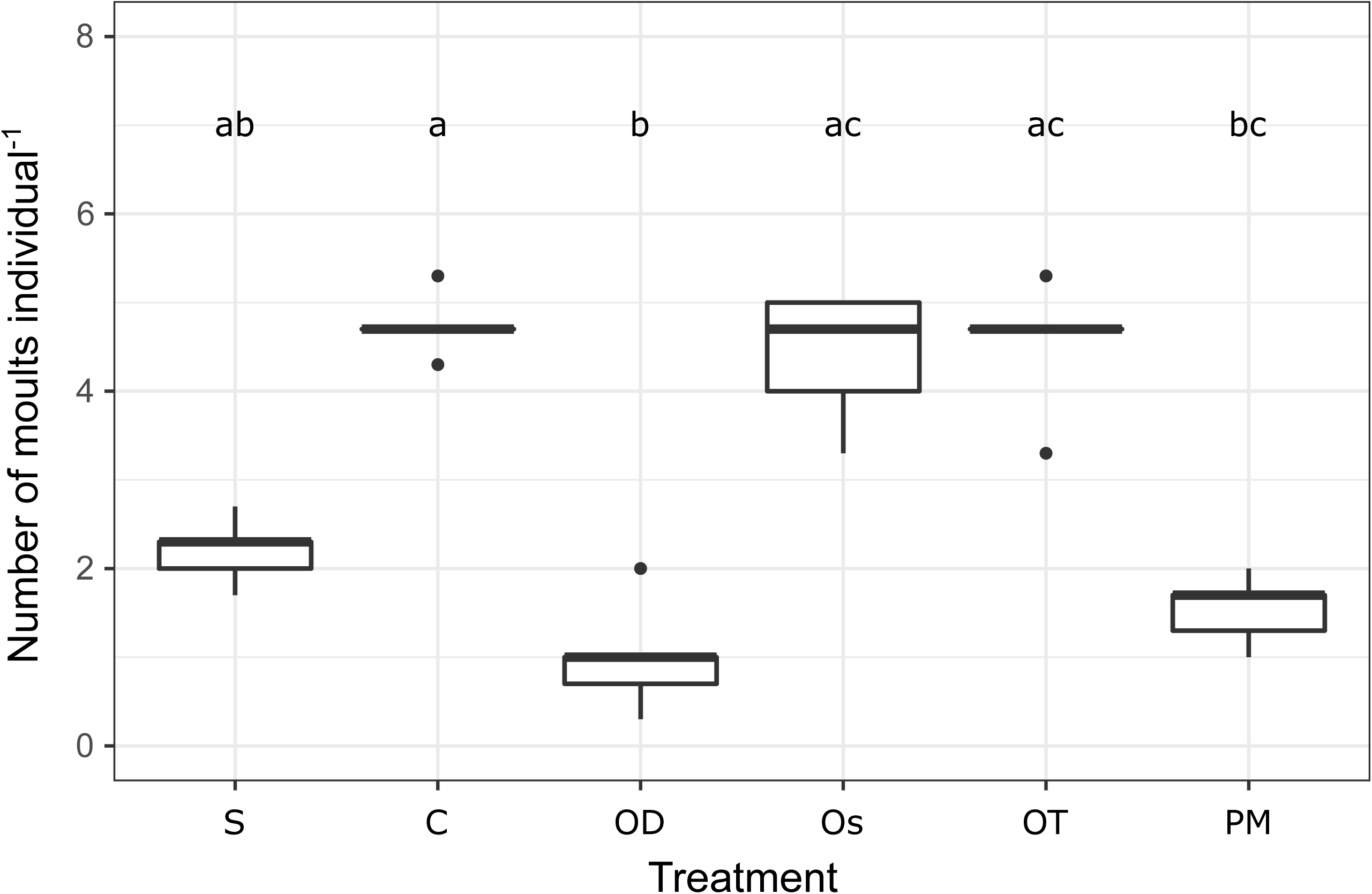
Cumulative number of moults (a proxy for growth) during the incubation period (11 days, *N*=5 per treatment). Letters indicate significant differences between treatments (p<0.05) and are based on Dunn’s post hoc test performed after a significant Kruskal-Wallis test. Treatments: Starvation treatment (S), the reference food *Cryptomonas* sp. (C), *Ochromonas danica* (OD), *Ochromonas* sp. (OS), *Ochromonas tuberculata* (OT), *Poterioochromonas malhamensis* (PM).

Consistent with the results on survival and growth, *D. longispina* only reached maturity on *Cryptomonas* sp., *Ochromonas sp*. and *O. tuberculata*, and not in the other treatments. Interestingly, the mean number of mature *D. longispina* (i.e., with eggs or embryos) at the time of termination of the experiment was significantly higher when feeding on *Ochromonas* sp. and *O. tuberculata* compared to the reference food *Cryptomonas* sp. (Kruskal-Wallis test: χ2 = 9.8207, df = 2, p = < 0.01, Table 4). Within these two chrysophyte treatments, almost every individual developed eggs or had juveniles at the time of termination.

**Table 4.** Mean (± SD) number of mature *D. longispina* (i.e., with eggs, embryos or juveniles) on the day of termination in the three treatments in which they were able to reach maturity. Analysis is based on Kruskal-Wallis test and subsequent Dunn’s post hoc test (*N*=5). Asterisks indicate significant differences compared to *Cryptomonas* sp. (p<0.05)

| **Treatment** | **No. of females with eggs / juveniles** |
| --- | --- |
| *Cryptomonas* sp. | $0.26 \pm 0.26$ |
| *Ochromonas* sp. | $0.90 \pm 0.20$ * |
| *Ochromonas tuberculata* | $0.94 \pm 0.12$ * |

## 4. Discussion

The feeding experiment clearly showed that dietary effects of mixotrophic chrysophytes can vary considerably even among taxonomically closely related species, from beneficial to deleterious consequences for *Daphnia*. Chrysophytes do not necessarily represent a suboptimal food source compared to cryptophytes. Thus, dietary value of freshwater mixotrophs cannot be predicted by their respective affiliation to major taxonomic groups. The fact that species with similar size, elemental stoichiometry, and fatty acid composition can largely differ in their dietary effects shows that other traits, such as toxicity, need to be put in focus of future research to understand the underlying mechanisms.

In the genera *Ochromonas* and *Poterioochromonas*, adverse effects on zooplankton were reported (Table 1). Toxic compounds produced by members of these genera were also detected (Reich & Spiegelstein, 1964; Halevy, Saliternik & Avivi, 1971), and the existing evidence suggest that toxins are intracellular substances (Table 1). Our results point out that the negative effects of chrysophytes on zooplankton cannot be generalised. Among the tested taxa, *O. danica* and *P. malhamensis* had clear negative dietary effects on zooplankton, but *O. tuberculata* and *Ochromonas* sp. proved to be adequate food largely comparable or, in case of first reproduction, even better than the reference diet *Cryptomonas* sp.

Especially *O. danica* suppressed zooplankton survival and growth. Feeding on this chrysophyte resulted in a severe drop (∼50%) of *D. longispina* survival in the first few days of the experiment, which suggests acute toxic effects. Although the identification of toxins was not part of the present study, the fact that the percentage of surviving *D. longispina* was significantly lower in the first 10 days than in the starvation treatment during the whole experimental duration suggests toxic diet effects (Lampert, 1981). It is highly unlikely that poor food quality alone would result in rapid mortality. Interestingly, following the elevated mortality in the initial phase, survival stabilised. This may be a sign of acclimatization to toxins, which may happen through physiological responses (Schwarzenberger *et al*., 2010) or changes in the gut microbiota (Macke *et al*., 2017). However, this was not coupled with growth, as indicated by the lack of moults. With the exception of one single individual, all juveniles died by the end of the 11-day experimental exposure before reaching maturity. Less acute, but still negative dietary effects were observed on *P. malhamensis*, with mass mortality occurring only in the second part of the experiment. This chrysophyte also suppressed growth (as indicated by the lack of moulting) and all *D. longispina* died during the incubation period without reaching maturity. These results are altogether consistent with the findings of Hiltunen, Barreiro & Hairston (2012) on *O. danica* and Zhang *et al*. (2011) on *P. malhamensis*. Both studies suggested toxic effects of these chrysophytes on their consumers, however, *O. danica* seemed to be more detrimental. This is in line with an earlier observation that *O. danica* contains higher cellular concentrations of toxins than *P. malhamensi*s (Reich & Spiegelstein, 1964). Differences in toxin levels may also well explain the less acute effect of *P. malhamensis* on *D. longispina* found in our study, but as this was similar to the starvation treatment, inadequate nutritional quality cannot be excluded either as the underlying mechanism. Additionally, we could observe a tendency of *P. malhamensis* to form sedimenting aggregates despite regular mixing. Hence, the possibility that an acute toxic effect of *P. malhamensis* was mitigated by lower food ingestion due to sedimentation cannot be excluded.

In contrast, the other two chrysophytes, *O. tuberculata* and *Ochromonas* sp. appeared to be adequate food sources, their dietary effects were largely comparable to the reference food *Cryptomonas* sp. On both of these treatments, only one single *D. longispina* died early in the experiment and all others reached maturity. Interestingly, *D. longispina* fed with these two *Ochromonas* strains started to reproduce earlier than on the reference food *Cryptomonas* sp.. The result for *O. tuberculata* is in clear contrast to previous findings on the species, which reported toxic effects and no reproduction of *Daphnia magna* due to high mortality (Katechakis *et al*., 2005). This is particularly intriguing because the same strain as in the present study was used.

Toxin production in ‘toxic’ algae often is strain-specific (Otsuka *et al*., 1999) and often depends on cellular physiological state and environmental cues (Watanabe & Oishi, 1985). The same holds for nutritional quality (Hessen, Færøvig & Andersen, 2002). In mixotrophs, cellular stoichiometry was found to be less dependent on light and nutrient conditions than in autotrophs, which may ensure more stable secondary production (Katechakis *et al*., 2005; Moorthi *et al*., 2017). However, nutritional mode can modulate dietary quality, and existing studies suggest that osmotrophic nutrition may enhance deleterious effects (Leeper & Porter, 1995; Hiltunen *et al*., 2012) and modulate fatty acid profiles (Boëchat *et al*., 2007). These altogether suggest that dietary quality of mixotrophs may dynamically vary through space and time. In the bioassay of the present study, all algae were cultivated under conditions inducing predominantly photo-autotrophic nutrition, and it is possible that the observed effects on *Daphnia* may change with cultivation conditions. However, the huge differences in dietary responses resulting from algae grown under standardised conditions suggest that species identity also plays a key role in determining the nutritional value of chrysophytes.

Given the somewhat different experimental conditions (lower P concentration in the medium, but higher frequency of glucose addition in the food quality experiment), linking the recorded stoichiometric and biochemical composition to the outcome of the feeding experiments has to be treated with caution. However, it is possible to draw conclusions on the nutritional value of algae and species-specific differences resulting from an experiment with standardised cultivation conditions. The very similar cellular C:P ratios suggest that no species-specific stoichiometric differences can be expected among these chrysophytes, given similar environmental conditions. All chrysophytes had C:P ratios (100-200) favourable for *Daphnia* growth (Hessen *et al*., 2013; Khattak *et al*., 2018). However, the fatty acid composition of these chrysophytes did not match the anticipated consumer growth response. *O. danica* had higher quantities of essential fatty acids, and PUFA in general, than the other two chrysophytes. This was not reflected in the outcome of the feeding bioassay, where it represented the poorest food source, and altogether suggests that the nutritional quality, as often assessed by fatty acids and PUFA in particular (Mariash *et al*., 2011; Galloway *et al*., 2014; Strandberg *et al*., 2015), may not be the main driver of the observed dietary effects of chrysophytes.

### Implications and future studies

In addition to grazing avoidance via colony formation (e.g., *Uroglenopsis* or *Dinobryon*) or scales, spines, and lorica (e.g. Vad et al., 2020), our results imply that species-specific differences in toxicity can also be important drivers of predator-prey interactions between mixotrophic chrysophytes and zooplankton. Although toxin production may be triggered by environmental cues or the presence of predators, we currently lack mechanistic understanding of toxin production in mixotrophs. To better understand the trophic effect of mixotrophic algae in aquatic food webs, we suggest three following avenues for future studies.

Firstly, toxicity seems to be a common trait in chrysophytes (Table 1), however, it may depend on physiological strategies, e.g., direct (osmotrophy) and indirect (phagotrophy) use of dissolved organic carbon can result in different consequences on consumers (Leeper & Porter, 1995; Hiltunen et al., 2012). Notably, the two species (*O. danica, P. malhamensis*) in our experiment which showed indications for toxicity were able to survive and grow in the dark, the others not (preliminary experiments, data not shown). This could point at fundamental differences in physiological strategies between these species. Future studies may specifically test for different physiological strategies as underlying mechanisms to shape the nutritional adequacy of mixotrophs for consumers.

Secondly, understanding the impact of mixotrophs on their consumers is critical in terms of environmental change. Human impacts and climate change are expected to increase the importance of mixotrophic protists in phytoplankton communities (Bergström et al., 2003; Wilken et al., 2018). In addition, physiological changes in mixotrophs, specifically a metabolic shift towards heterotrophy due to increasing water temperatures are expected (Wilken et al., 2013). This is particularly important if physiological changes induce changes in food quality.

Thirdly, existing evidence on the nutritional adequacy of chrysophytes originates mostly from controlled experimental studies. These, together with a limited number of observational studies from natural lakes suggest a potentially influential role in energy transfer towards higher trophic levels (Bertoni *et al*., 2002; Vad *et al*., 2020). Community-level studies involving more realistic food webs may prove important to better understand the relevance of increased dominance of chrysophytes on higher trophic levels.

## Acknowledgements

This project has received funding from the European Union’s Horizon 2020 research and innovation programme under the Marie Skłodowska-Curie Grant agreement No. 658400. CFV acknowledges further support by the GINOP 2.3.2.-15-2016-00057 and NKFIH-872 projects, and the AQUACOSM-plus project of the European Union’s Horizon 2020 research and innovation programme under grant agreement No. 871081. RP and RF acknowledge support by FWF (I 3311-B25) and by the H2020 research and innovation programme under grant agreement No. 731065 (AQUACOSM). The authors wish to thank Achim Weigert for performing the elemental analyses and Samuel-Karl Kämmer for the fatty acid analyses, and three anonymous reviewers for their constructive feedback and comments.

## Data Availability Statement

The data that support the findings of this study are available from the corresponding author upon reasonable request.

## Author contributions

CFV, CS, RF and RP conceived the study. CS, RF and CFV performed the laboratory experiments. MJK carried out the biochemical analyses. CFV and CS analysed the data. CFV and CS wrote the first draft after which all authors provided critical feedback on the manuscript.

## References

Ahlgren G., Lundstedt L., Brett M. & Forsberg C. (1990). Lipid composition and food quality of some freshwater phytoplankton for cladoceran zooplankters. Journal of Plankton Research 12, 809–818. https://doi.org/10.1093/plankt/12.4.809

Anderson R., Jürgens K. & Hansen P.J. (2017). Mixotrophic Phytoflagellate Bacterivory Field Measurements Strongly Biased by Standard Approaches: A Case Study. Frontiers in Microbiology 8. https://doi.org/10.3389/fmicb.2017.01398

Beisner B.E., Grossart H.-P. & Gasol J.M. (2019). A guide to methods for estimating phago-mixotrophy in nanophytoplankton. Journal of Plankton Research 41, 77–89. https://doi.org/10.1093/plankt/fbz008

Bertoni R., Callieri C. & Corno G. (2002). The mixotrophic flagellates as key organisms from DOC to Daphnia in an oligotrophic alpine lake. Internationale Vereinigung fur Theoretische und Angewandte Limnologie Verhandlungen 28, 392–395

Bird D.F. & Kalff J. (1987). Algal phagotrophy: regulating factors and importance relative to photosynthesis in Dinobryon(Chrysophyceae). Limnology and Oceanography 32, 277– 284

Boëchat I.G., Weithoff G., Krüger A., Gücker B. & Adrian R. (2007). A biochemical explanation for the success of mixotrophy in the flagellate Ochromonas sp. Limnology and Oceanography 52, 1624–1632. https://doi.org/10.4319/lo.2007.52.4.1624

Boenigk J. & Stadler P. (2004). Potential toxicity of chrysophytes affiliated with Poterioochromonas and related ‘Spumella-like’flagellates. Journal of Plankton Research 26, 1507–1514. https://doi.org/10.1093/plankt/fbh139

Boenigk J., Stadler P., Wiedlroither A. & Hahn M.W. (2004). Strain-Specific Differences in the Grazing Sensitivities of Closely Related Ultramicrobacteria Affiliated with the Polynucleobacter Cluster. Applied and Environmental Microbiology 70, 5787–5793. https://doi.org/10.1128/AEM.70.10.5787-5793.2004

Boxhorn J.E., Holen D.A. & Boraas M.E. (1998). Toxicity of the Chrysophyte flagellate Poterioochromonas malhamensis to the rotifer Brachionus angularis. Hydrobiologia 387, 283–287. https://doi.org/10.1023/A:1017030919264

Brett M. & Müller-Navarra D. (1997). The role of highly unsaturated fatty acids in aquatic foodweb processes. Freshwater Biology 38, 483–499. https://doi.org/10.1046/j.1365-2427.1997.00220.x

Burkholder J.M., Glibert P.M. & Skelton H.M. (2008). Mixotrophy, a major mode of nutrition for harmful algal species in eutrophic waters. Harmful Algae 8, 77–93. https://doi.org/10.1016/j.hal.2008.08.010

Carpenter S.R. & Kitchell J.F. (1996). The Trophic Cascade in Lakes. Cambridge University Press.

Domaizon I., Viboud S. & Fontvieille D. (2003). Taxon-specific and seasonal variations in flagellates grazing on heterotrophic bacteria in the oligotrophic Lake Annecy – importance of mixotrophy. FEMS Microbiology Ecology 46, 317–329. https://doi.org/10.1016/S0168-6496(03)00248-4

van Donk E., Cerbin S., Wilken S., Helmsing N.R., Ptacnik R. & Verschoor A.M. (2009). The effect of a mixotrophic chrysophyte on toxic and colony-forming cyanobacteria. Freshwater Biology 54, 1843–1855. https://doi.org/10.1111/j.1365-2427.2009.02227.x

von Elert E. & Stampfl P. (2000). Food quality for Eudiaptomus gracilis: the importance of particular highly unsaturated fatty acids. Freshwater Biology 45, 189–200. https://doi.org/10.1046/j.1365-2427.2000.00671.x

Fischer R., Giebel H.-A., Hillebrand H. & Ptacnik R. (2017). Importance of mixotrophic bacterivory can be predicted by light and loss rates. Oikos 126, 713–722. https://doi.org/10.1111/oik.03539

Flynn K.J. & Mitra A. (2009). Building the “perfect beast”: modelling mixotrophic plankton. Journal of Plankton Research 31, 965–992. https://doi.org/10.1093/plankt/fbp044

Flynn K.J., Mitra A., Glibert P.M. & Burkholder J.M. (2018). Mixotrophy in Harmful Algal Blooms: By Whom, on Whom, When, Why, and What Next. In: Global Ecology and Oceanography of Harmful Algal Blooms. Ecological Studies, (Eds P.M. Glibert, E. Berdalet, M.A. Burford, G.C. Pitcher & M. Zhou), pp. 113–132. Springer International Publishing, Cham.

Galloway A.W.E., Taipale S.J., Hiltunen M., Peltomaa E., Strandberg U., Brett M.T., et al. (2014). Diet-specific biomarkers show that high-quality phytoplankton fuels herbivorous zooplankton in large boreal lakes. Freshwater Biology 59, 1902–1915. https://doi.org/10.1111/fwb.12394

Hahn M.W. & Höfle M.G. (1998). Grazing Pressure by a Bacterivorous Flagellate Reverses the Relative Abundance of Comamonas acidovoransPX54 and Vibrio Strain CB5 in Chemostat Cocultures. Applied and Environmental Microbiology 64, 1910–1918. https://doi.org/10.1128/AEM.64.5.1910-1918.1998

Halevy S., Saliternik R. & Avivi L. (1971). Isolation of rhodamine-positive toxins from Ochromonas and other algae. International Journal of Biochemistry 2, 185–192. https://doi.org/10.1016/0020-711X(71)90211-4

Hansen H.P. & Koroleff F. (1999). Determination of nutrients. In: Methods of Seawater Analysis, 3rd edn. (Eds K. Grasshoff, K. Kremling & M. Ehrhardt), pp. 159–228. Wiley-VCH, Weinheim.

Hartmann M., Grob C., Tarran G.A., Martin A.P., Burkill P.H., Scanlan D.J., et al. (2012). Mixotrophic basis of Atlantic oligotrophic ecosystems. Proceedings of the National Academy of Sciences of the United States of America 109, 5756–5760. https://doi.org/10.1073/pnas.1118179109

Heissenberger M., Watzke J. & Kainz M.J. (2010). Effect of nutrition on fatty acid profiles of riverine, lacustrine, and aquaculture-raised salmonids of pre-alpine habitats. Hydrobiologia 650, 243–254. https://doi.org/10.1007/s10750-010-0266-z

Hessen D.O., Elser J.J., Sterner R.W. & Urabe J. (2013). Ecological stoichiometry: An elementary approach using basic principles. Limnology and Oceanography 58, 2219– 2236. https://doi.org/10.4319/lo.2013.58.6.2219

Hessen D.O., Færøvig P.J. & Andersen T. (2002). Light, Nutrients, and P:c Ratios in Algae: Grazer Performance Related to Food Quality and Quantity. Ecology 83, 1886–1898. https://doi.org/10.1890/0012-9658(2002)083[1886:LNAPCR]2.0.CO;2

Hessen D.O. & Rukke N.A. (2000). The costs of moulting in Daphnia; mineral regulation of carbon budgets. Freshwater Biology 45, 169–178. https://doi.org/10.1046/j.1365-2427.2000.00670.x

Hillebrand H., Dürselen C.-D., Kirschtel D., Pollingher U. & Zohary T. (1999). Biovolume Calculation for Pelagic and Benthic Microalgae. Journal of Phycology 35, 403–424. https://doi.org/10.1046/j.1529-8817.1999.3520403.x

Hiltunen T., Barreiro A. & Hairston N.G. (2012). Mixotrophy and the toxicity of Ochromonas in pelagic food webs. Freshwater Biology 57, 2262–2271. https://doi.org/10.1111/fwb.12000

Isaksson A., Bergström A.-K., Blomqvist P. & Jansson M. (1999). Bacterial grazing by phagotrophic phytoflagellates in a deep humic lake in northern Sweden. Journal of Plankton Research 21, 247–268

Kamjunke N., Henrichs T. & Gaedke U. (2007). Phosphorus gain by bacterivory promotes the mixotrophic flagellate Dinobryon spp. during re-oligotrophication. Journal of Plankton Research 29, 39–46. https://doi.org/10.1093/plankt/fbl054

Kamjunke N., Köhler B., Wannicke N. & Tittel J. (2008). Algae as Competitors for Glucose with Heterotrophic Bacteria. Journal of Phycology 44, 616–623. https://doi.org/10.1111/j.1529-8817.2008.00520.x

Katechakis A., Haseneder T., Kling R. & Stibor H. (2005). Mixotrophic versus photoautotrophic specialist algae as food for zooplankton: The light: nutrient hypothesis might not hold for mixotrophs. Limnology and Oceanography 50, 1290– 1299. https://doi.org/10.4319/lo.2005.50.4.1290

Katechakis A. & Stibor H. (2006). The mixotroph Ochromonas tuberculata may invade and suppress specialist phago- and phototroph plankton communities depending on nutrient conditions. Oecologia 148, 692–701. https://doi.org/10.1007/s00442-006-0413-4

Keller M.D., Shapiro L.P., Haugen E.M., Cucci T.L., Sherr E.B. & Sherr B.F. (1994). Phagotrophy of fluorescently labeled bacteria by an oceanic phytoplankter. Microbial Ecology 28, 39–52. https://doi.org/10.1007/BF00170246

Khattak H.K., Prater C., Wagner N.D. & Frost P.C. (2018). The threshold elemental ratio of carbon and phosphorus of Daphnia magna and its connection to animal growth. Scientific Reports 8, 9673. https://doi.org/10.1038/s41598-018-27758-7

Kies L. (1967). über Zellteilung und Zygotenbildung bei Roya obtusa (Breb.) West et West. Mitteilungen des Staatsinstituts für Allgemeine Botanik Hamburg 12, 35–42

Lampert W. (1981). Inhibitory and Toxic Effects of Blue-green Algae on Daphnia. Internationale Revue der gesamten Hydrobiologie und Hydrographie 66, 285–298. https://doi.org/10.1002/iroh.19810660302

Leeper D.A. & Porter K.G. (1995). Toxicity of the mixotrophic chrysophyte Poterioochromonas malhamensis to the cladoceran Daphnia ambigua. Archiv für Hydrobiologie 134, 207–222

Leles S.G., Mitra A., Flynn K.J., Tillmann U., Stoecker D., Jeong H.J., et al. (2019). Sampling bias misrepresents the biogeographical significance of constitutive mixotrophs across global oceans. Global Ecology and Biogeography 28, 418–428. https://doi.org/10.1111/geb.12853

Macke E., Callens M., De Meester L. & Decaestecker E. (2017). Host-genotype dependent gut microbiota drives zooplankton tolerance to toxic cyanobacteria. Nature Communications 8, 1608. https://doi.org/10.1038/s41467-017-01714-x

Mariash H.L., Cazzanelli M., Kainz M.J. & Rautio M. (2011). Food sources and lipid retention of zooplankton in subarctic ponds. Freshwater Biology 56, 1850–1862. https://doi.org/10.1111/j.1365-2427.2011.02625.x

Martin-Creuzburg D., Elert E. von & Hoffmann K.H. (2008). Nutritional constraints at the cyanobacteria—Daphnia magna interface: The role of sterols. Limnology and Oceanography 53, 456–468. https://doi.org/10.4319/lo.2008.53.2.0456

Mitra A., Flynn K.J., Burkholder J.M., Berge T., Calbet A., Raven J.A., et al. (2014). The role of mixotrophic protists in the biological carbon pump. Biogeosciences 11, 995–1005. https://doi.org/10.5194/bg-11-995-2014

Moorthi S.D., Ptacnik R., Sanders R.W., Fischer R., Busch M. & Hillebrand H. (2017). The functional role of planktonic mixotrophs in altering seston stoichiometry. Aquatic Microbial Ecology 79, 235–245. https://doi.org/10.3354/ame01832

Müller-Navarra D. & Lampert W. (1996). Seasonal patterns of food limitation in Daphnia galeata: separating food quantity and food quality effects. Journal of Plankton Research 18, 1137–1157. https://doi.org/10.1093/plankt/18.7.1137

Nygaard K. & Tobiesen A. (1993). Bacterivory in algae: A survival strategy during nutrient limitation. Limnology and Oceanography 38, 273–279. https://doi.org/10.4319/lo.1993.38.2.0273

Otsuka S., Suda S., Li R., Watanabe M., Oyaizu H., Matsumoto S., et al. (1999). Phylogenetic relationships between toxic and non-toxic strains of the genus Microcystis based on 16S to 23S internal transcribed spacer sequence. FEMS Microbiology Letters 172, 15– 21. https://doi.org/10.1111/j.1574-6968.1999.tb13443.x

Pålsson C. & Granéli W. (2004). Nutrient limitation of autotrophic and mixotrophic phytoplankton in a temperate and tropical humic lake gradient. Journal of Plankton Research 26, 1005–1014. https://doi.org/10.1093/plankt/fbh089

Ptacnik R., Gomes A., Royer S.-J., Berger S.A., Calbet A., Nejstgaard J.C., et al. (2016). A light-induced shortcut in the planktonic microbial loop. Scientific Reports 6, srep29286. https://doi.org/10.1038/srep29286

Ptacnik R., Lepistö L., Willén E., Brettum P., Andersen T., Rekolainen S., et al. (2008). Quantitative responses of lake phytoplankton to eutrophication in Northern Europe. Aquatic Ecology 42, 227–236. https://doi.org/10.1007/s10452-008-9181-z

R Core Team (2020). R: A Language and Environment for Statistical Computing. R Foundation for Statistical Computing, Vienna, Austria.

Reich K. & Spiegelstein M. (1964). Fishtoxins in Ochromonas (chrysomonadina). Israel Journal of Zoology 13, 141–141. https://doi.org/10.1080/00212210.1964.10688196

Sanders R.W. & Porter K.G. (1988). Phagotrophic Phytoflagellates. In: Advances in Microbial Ecology. Advances in Microbial Ecology, (Ed. K.C. Marshall), pp. 167–192. Springer US, Boston, MA.

Schwarzenberger A., Zitt A., Kroth P., Mueller S. & Von Elert E. (2010). Gene expression and activity of digestive proteases in Daphnia: effects of cyanobacterial protease inhibitors. BMC Physiology 10, 6. https://doi.org/10.1186/1472-6793-10-6

Solórzano L. & Sharp J.H. (1980). Determination of total dissolved phosphorus and particulate phosphorus in natural waters. Limnology and Oceanography 25, 754–758. https://doi.org/10.4319/lo.1980.25.4.0754

Sommer U. & Stibor H. (2002). Copepoda – Cladocera – Tunicata: The role of three major mesozooplankton groups in pelagic food webs. Ecological Research 17, 161–174. https://doi.org/10.1046/j.1440-1703.2002.00476.x

Sterner R.W. & Elser J.J. (2002). Ecological Stoichiometry: The Biology of Elements from Molecules to the Biosphere. Princeton University Press.

Sterner R.W. & Hessen D.O. (1994). Algal Nutrient Limitation and the Nutrition of Aquatic Herbivores. Annual Review of Ecology and Systematics 25, 1–29. https://doi.org/10.1146/annurev.es.25.110194.000245

Sterner R.W. & Schulz K.L. (1998). Zooplankton nutrition: recent progress and a reality check. Aquatic Ecology 32, 261–279. https://doi.org/10.1023/A:1009949400573

Stoecker D.K., Hansen P.J., Caron D.A. & Mitra A. (2017). Mixotrophy in the Marine Plankton. Annual Review of Marine Science 9, 311–335. https://doi.org/10.1146/annurev-marine-010816-060617

Strandberg U., Taipale S.J., Hiltunen M., Galloway A.W.E., Brett M.T. & Kankaala P. (2015). Inferring phytoplankton community composition with a fatty acid mixing model. Ecosphere 6, art16. https://doi.org/10.1890/ES14-00382.1

Tang H., Zhu S., Wang N., Xu Z., Huang J., Gu L., et al. (2020). The inhibitory effect of mixotrophic Ochromonas gloeopara on the survival and reproduction of Daphnia similoides sinensis. Environmental Science and Pollution Research 27, 29068–29074. https://doi.org/10.1007/s11356-020-09291-1

Tranvik L.J., Porter K.G. & Sieburth J.McN. (1989). Occurrence of bacterivory in Cryptomonas, a common freshwater phytoplankter. Oecologia 78, 473–476. https://doi.org/10.1007/BF00378736

Unrein F., Gasol J.M., Not F., Forn I. & Massana R. (2014). Mixotrophic haptophytes are key bacterial grazers in oligotrophic coastal waters. The ISME Journal 8, 164–176. https://doi.org/10.1038/ismej.2013.132

Urabe J., Gurung T.B. & Yoshida T. (1999). Effects of phosphorus supply on phagotrophy by the mixotrophic alga Uroglena americana (Chrysophyceae). Aquatic Microbial Ecology 18, 77–83. https://doi.org/10.3354/ame018077

Vad C.F., Schneider C., Lukić D., Horváth Z., Kainz M.J., Stibor H., et al. (2020). Grazing resistance and poor food quality of a widespread mixotroph impair zooplankton secondary production. Oecologia 193, 489–502. https://doi.org/10.1007/s00442-020-04677-x

Vadstein O., Jensen A., Olsen Y. & Reinertsen H. (1988). Growth and phosphorus status of limnetic phytoplankton and bacteria. Limnology and Oceanography 33, 489–503

Ward B.A. & Follows M.J. (2016). Marine mixotrophy increases trophic transfer efficiency, mean organism size, and vertical carbon flux. Proceedings of the National Academy of Sciences 113, 2958–2963. https://doi.org/10.1073/pnas.1517118113

Watanabe M.F. & Oishi S. (1985). Effects of Environmental Factors on Toxicity of a Cyanobacterium (Microcystis aeruginosa) under Culture Conditions. Applied and Environmental Microbiology 49, 1342–1344

Watson S.B., McCauley E. & Downing J.A. (1997). Patterns in phytoplankton taxonomic composition across temperate lakes of differing nutrient status. Limnology and Oceanography 42, 487–495. https://doi.org/10.4319/lo.1997.42.3.0487

Watson S.B., Whitton B.A., Higgins S.N., Paerl H.W., Brooks B.W. & Wehr J.D. (2015). Chapter 20 - Harmful Algal Blooms. In: Freshwater Algae of North America, 2nd Edition. Aquatic Ecology, (Eds J.D. Wehr, R.G. Sheath & J.P. Kociolek), pp. 873–920. Academic Press, Boston.

Zhang X., Hu H.-Y., Warming T.P. & Christoffersen K.S. (2011). Life History Response of Daphnia magna to a Mixotrophic Golden Alga, Poterioochromonas sp., at Different Food Levels. Bulletin of Environmental Contamination and Toxicology 87, 117–123. https://doi.org/10.1007/s00128-011-0328-6

Zubkov M.V. & Tarran G.A. (2008). High bacterivory by the smallest phytoplankton in the North Atlantic Ocean. Nature 455, 224–226. https://doi.org/10.1038/nature07236

