## Supplementary material for "From adverse to beneficial – contrasting dietary effects of freshwater mixotrophs on zooplankton": Table S1

**Supporting information**

Table S1 – Mass fractions (µg mg dry weight^-1^; mean ± SD) of individual fatty acids (FA) in chrysophytes and *Cryptomonas* sp.. Essential fatty acids are highlighted in bold. (MUFA – monounsaturated fatty acids, PUFA – polyunsaturated fatty acids, SAFA – saturated fatty acids).

|  |  | *Cryptomonas* sp. |  | *O. danica* |  | *O. tuberculata* |  | *P. malhamensis* |
| --- | --- | --- | --- | --- | --- | --- | --- | --- |
| SAFA |  |  |  |  |  |  |  |  |
| 14:0 |  | 5.3 ±1.5 |  | 9.7 ± 0.9 |  | 5.6 ± 1.6 |  | 4.8 ± 0.8 |
| 15:0 |  | 0.7 ±0.1 |  | 0.7 ± 0.1 |  | 0.4 ± 0.1 |  | 1.0 ± 0.0 |
| 16:0 |  | 15.5 ±3.6 |  | 31.8 ± 4.8 |  | 20.6 ± 8.5 |  | 24.3 ± 3.3 |
| 17:0 |  | 0.5 ±0.2 |  | 0.1 ± 0.0 |  | 0.1 ± 0.1 |  | 0.1 ±0.0 |
| 18:0 |  | 5.4 ±1.9 |  | 17.3 ± 2.7 |  | 12.0 ± 5.1 |  | 13.1 ± 2.1 |
| 20:0 |  | 0.0 ±0.0 |  | 0.0 ± 0.0 |  | 0.0 ± 0.0 |  | 0.0 ± 0.0 |
| MUFA |  |  |  |  |  |  |  |  |
| 16.1n-9 |  | 0.9 ±0.4 |  | 1.7 ± 0.9 |  | 0.9 ± 0.3 |  | 4.5 ± 0.3 |
| 16.1n-7 |  | 4.6 ±2.1 |  | 1.3 ± 0.3 |  | 3.5 ± 1.4 |  | 0.6 ± 0.1 |
| 18.1n-12 |  | 0.0 ±0.0 |  | 0.0 ± 0.0 |  | 0.0 ± 0.0 |  | 0.0 ± 0.0 |
| 18.1n-9c |  | 3.1 ±0.5 |  | 3.4 ± 0.7 |  | 1.1 ± 0.3 |  | 1.3 ± 0.0 |
| 18.1n-7 |  | 5.7 ±1.2 |  | 8.1 ± 1.2 |  | 3.0 ± 1.1 |  | 5.4 ± 1.5 |
| PUFA |  |  |  |  |  |  |  |  |
| **18.2n-6** |  | 6.7 ±1.2 |  | 8.9 ± 1.2 |  | 2.4 ± 1.0 |  | 1.5 ± 0.4 |
| 18.3n-6 |  | 0.3 ±0.1 |  | 0.8 ± 0.1 |  | 0.5 ± 0.1 |  | 0.2 ± 0.1 |
| **18.3n-3** |  | 9.7 ±0.4 |  | 8.1 ± 2.1 |  | 1.2 ± 0.4 |  | 0.7 ± 0.3 |
| 18.4n-3 |  | 5.0 ±0.8 |  | 1.9 ± 0.1 |  | 0.9 ± 0.3 |  | 0.2 ± 0.1 |
| 20.3n-6 |  | 0.2 ±0.4 |  | 1.8 ± 0.1 |  | 0.0 ± 0.0 |  | 0.0 ± 0.0 |
| **20.4n-6** |  | 0.9 ±0.8 |  | 4.0 ± 0.6 |  | 0.0 ± 0.0 |  | 0.0 ± 0.0 |
| **20.5n-3** |  | 6.8 ±0.7 |  | 2.1 ± 0.3 |  | 0.0 ± 0.0 |  | 0.1 ± 0.1 |
| 22.3n-3 |  | 0.7 ±0.8 |  | 3.6 ± 0.4 |  | 1.9 ± 0.9 |  | 1.9 ± 0.3 |
| 22.5n-3 |  | 0.0 ±0.0 |  | 0.0 ± 0.0 |  | 0.0 ± 0.0 |  | 0.2 ± 0.3 |
| **22.6n-3** |  | 0.8 ±0.2 |  | 0.5 ± 0.1 |  | 0.0 ± 0.1 |  | 0.3 ± 0.1 |
